# Neural reinstatement of features in audiovisual working memory indicates object-based retrieval

**DOI:** 10.64898/2026.02.06.704386

**Authors:** Ceren Arslan, Daniel Schneider, Stephan Getzmann, Edmund Wascher, Laura-Isabelle Klatt

**Affiliations:** Leibniz Research Centre for Working Environment and Human Factors; Donders Institute for Brain, Cognition and Behaviour, Radboud University; Institute of Medical Psychology, Goethe University Frankfurt

**Keywords:** working memory, multisensory processing, feature-based attention, object-based attention, EEG, multivariate pattern analysis

## Abstract

Selective attention allows us to prioritize or retrieve task-relevant features and items from working memory. However, previous work has largely relied on unisensory paradigms, leaving open the question of how attentional mechanisms act on audiovisual working memory representations. Here, using an EEG-based audiovisual delayed-match-to-sample task, we investigate whether attention to audiovisual working memory contents operates on the level of individual unisensory features or bound cross-modal objects. On each trial, participants memorized an audiovisual item. At test, they were randomly presented with either an auditory, visual, or an audiovisual probe stimulus and indicated whether the latter matched their working memory content. Compared with recalling the entire audiovisual object, recall of unisensory visual or auditory features resulted in poorer behavioral performance and elevated midfrontal theta power. Multivariate pattern analyses (MVPA) showed that task-irrelevant feature representations were retrieved in both auditory and visual probe trials, consistent with object-based retrieval. Moreover, these results shed light on the representational structure underlying cross-modal feature storage in working memory, suggesting that audiovisual features are stored as bound objects and that attentional selection of individual object features in one modality spreads to cross-modal object features in another modality. In task conditions where such incidental recall of task-irrelevant features is detrimental to task performance, this places greater demands on cognitive control mechanisms.

## 1. Introduction

Working memory allows us to temporarily store a limited amount of information (Baddeley, 1992; Oberauer et al., 2016), making attentional selection of behaviorally relevant over irrelevant information essential for efficient memory performance (Baddeley, 2012; Gazzaley & Nobre, 2012). This selection not only occurs at the level of encoding, but also after information has already entered working memory (Myers et al., 2017). Accordingly, an ample body of work has shown that retro-cues directing attention to a task-relevant object incur behavioral benefits (reviewed by Souza & Oberauer, 2016). Similar retro-cue benefits have been observed when the cue indicates a task-relevant feature dimension (Hajonides, Van Ede, et al., 2020; Heuer et al., 2016; Heuer & Schubö, 2016; Park et al., 2017). Notably, in line with an object-based account of attention (Desimone & Duncan, 1995), a range of studies corroborates the notion that attention to individual object features will spread to other features of that same object in working memory. For instance, incidental cues presented during the maintenance period, which are uninformative about which item will be probed, have been shown to enhance memory for the uncued feature of the same visual object (Zokaei, Manohar, et al., 2014; Zokaei, Ning, et al., 2014). On a similar note, a range of behavioral studies in visual (Gao et al., 2011; Hyun et al., 2009; Jiang et al., 2016; Logie et al., 2011; Saltzmann et al., 2023; Shen et al., 2013), auditory (Fischer et al., 2024.; Joseph et al., 2015), and audiovisual (Arslan et al., 2025a; 2025b) working memory has shown that attending to one object-feature (e.g., the frequency of a sound) leads to the involuntary maintenance of another, task-irrelevant object feature (e.g., the sound’s location). Critically, a study by Printzlau et al. (2022) recently provided complementary neural evidence showing that presenting an uninformative color cue during maintenance reinstates the neural representation of its associated orientation, albeit the latter was task-irrelevant at the time of cue presentation. Together, these findings indicate that object-based attention can incidentally strengthen the representation of an object’s associated features. However, it remains unresolved whether similar mechanisms govern attentional selection within audiovisual working memory. Hence, this study asks whether selectively retrieving a single feature of a cross-modal object results in the reactivation of task-irrelevant features in another modality.

To date, incidental reactivation of associated features has primarily been examined from a multisensory perspective in the episodic memory literature. For example, Nyberg et al. (2000) presented participants with visual words, either alone or paired with complex sounds, during encoding, and subsequently tested visual word recognition. During retrieval, auditory cortex and medial temporal lobe regions that were active during the encoding of word–sound pairs were reactivated, despite auditory information being task-irrelevant (see Wheeler et al. (2000), for converging evidence of sensory-specific reinstatement in an audiovisual context). Similar findings have been reported in the olfactory domain, where images previously paired with odors reactivated not only memory-related regions but also primary olfactory cortex during retrieval (Gottfried et al., 2004). Together, these studies indicate that modality-specific sensory regions engaged during multisensory encoding can be spontaneously reactivated by unimodal targets from a different modality during retrieval, even when the associated sensory information is task-irrelevant. However, whether such cross-modal reinstatement occurs in audiovisual working memory has not yet been investigated.

While the reactivation of an object’s associated features can enhance memory performance when those features remain task-relevant (e.g., Printzlau et al., 2022), it may be detrimental to the performance when they are irrelevant. For example, using pre-cues, a study by Joseph et al. (2015) showed that recalling a single feature from a bound auditory object impairs performance relative to remembering the entire sound, an effect described as the feature extraction cost. Although this cost was interpreted as reflecting the ‘breaking up’ of a bound object representation, the authors noted that it may also arise from interference of task-irrelevant features during encoding, maintenance, or recall. If the interference occurred at recall, this would suggest that feature probes reinstated associated but irrelevant features of memory items. In this sense, feature extraction costs may reflect a conflict between object-based storage and the task demand to attend to a single feature selectively. When such competition between object-based and feature-based attentional selection emerges, additional regulatory mechanisms may be required to resolve this conflict. Cognitive control, which refers to the goal-directed regulation of behavior in the presence of competing or automatic processes (Cavanagh et al., 2012; Cavanagh & Frank, 2014; Cohen, 2014), may therefore be engaged when feature-based prioritization and spontaneous reactivation of object-based representations create a conflict.

Here, we combine behavioral methods and EEG to assess whether attentional selection during recall operates in a feature- or object-based fashion in audiovisual working memory. To investigate this question, participants performed an audiovisual delayed match-to-sample task. Critically, while the memory item always consisted of a visual and an auditory feature, the probe could be either visual, auditory, or audiovisual. This design allowed us to test whether attending to only one feature of an audiovisual object at recall automatically spreads attention to the behaviorally irrelevant feature, consistent with object-based accounts. Specifically, by randomly varying the task-relevant dimensions at recall, we aimed to test whether incidental reactivation of cross-modal object features in neural activity emerges in multisensory working memory. If attentional selection is primarily object-based, we expect behavioral costs associated with feature-selective (relative to whole-object) recall as well as the neural reinstatement of irrelevant features upon unimodal probe presentation. We further expect increased demands on cognitive control to mitigate the conflict between object-based selection and feature-based task demands in unimodal probe trials. However, in the absence of object-based selection (i.e., if attention operates on segregated modality-specific representations), we would expect no behavioral cost and less reliable (or a lack of) decoding of task-irrelevant features. More broadly, the present findings also shed light on the representational structure of audiovisual working memory contents. That is, if task-irrelevant features are incidentally reactivated upon unisensory probe presentation, this implies that they were encoded and stored as bound object representations.

## 2. Materials and Method

### 2.1 Ethics Statement

This study received approval from the Ethics Committee of the Leibniz Research Centre for Working Environment and Human Factors and was carried out in accordance with the principles of the Declaration of Helsinki. All participants provided written informed consent and were compensated at either 12 euros per hour or with course credits.

### 2.2 Participants

A total of 42 healthy young adults participated in the experiment. In total, five participants were excluded. This includes two exclusions due to technical EEG issues, one participant who withdrew from the study, and two participants with extremely noisy data (rejected trials > 25% of all trials). This leaves a total of 37 datasets that entered the main analysis (22 female, 15 male; 4 left-handed), with a mean age of 23.97 years (*SD* = 3.55, range: 19–35).

Participants’ hearing acuity was assessed using pure-tone audiometry (Oscilla USB 330; Inmedico, Lystrup, Denmark) across eleven frequencies ranging from 125 Hz to 8 kHz. All participants demonstrated normal hearing (≤25 dB) across frequencies from 125 Hz to 3 kHz. Eight participants showed slightly elevated thresholds (30–40 dB) for frequencies ≥3 kHz. Visual acuity was measured using Landolt C optotypes at a distance of 1.5 meters, with scores calculated via logarithmic averaging (Bach & Kommerell, 1998). All participants exhibited good visual acuity (*M* = 1.35, *SD* = 1.26, range = 0.75–1.88).

### 2.3 Experimental setup and stimuli

The experiment took place in a soundproof, dimly lit room measuring 5.0 × 3.3 × 2.4 meters. Stimuli were presented using E-Prime software (version 3.0, Psychology Software Tools, Pittsburgh, PA). An AudioFile Stimulus Processor was used to synchronize the auditory stimuli with EEG triggers (Cambridge Research Systems, Rochester, UK). Visual stimuli were displayed on a centrally positioned 49-inch monitor with a resolution of 5120 × 1440 pixels and a 100 Hz refresh rate (Samsung, Seoul, South Korea). A full-range loudspeaker (SC 55.9–8 Ohm; Visaton, Haan, Germany) was centrally positioned below the screen at a height of ∼1.3 meters. Participants were seated 1.5 meters from the screen.

Visual memory items were eight sine-wave gratings with angles ranging from 11.25° to 168.75° (11.25°, 33.75°, 56.25°, 78.75°, 101.25°, 123.75°, 146.25°, 168.75°) presented at a 5% contrast on a gray screen (RGB = 150, 150, 150). The size of the gratings was 5.5° with 0.62 cycles per degree. The visual impulse was a white circle with a size of 9.5° (RGB = 255, 255, 255). Finally, as a visual filler item, a grey circle with a diameter of 5.5° was generated (RGB = 160, 160, 160). All visual stimuli were created using a custom-made MATLAB script (R2022a).

Auditory memory items consisted of eight harmonic tones with base frequencies of 270, 381, 540, 763, 1080, 1527, 2159, and 3054 Hz, created using Cool Edit 2000 1.1 (Syntrillium Software Corporation, Phoenix, USA). Each tone included five harmonics: the fundamental frequency at full amplitude (100%), followed by the second at 40%, the third at 30%, the fourth at 20%, and the fifth at 10%. A 10 ms fade-in and fade-out was applied to each tone, and all were sampled at 44,100 Hz. The auditory impulse was a normalized complex tone consisting of all memory item tones. Finally, as an auditory filler, pink noise was generated with an amplitude of 0.5 using the free software Audacity (version 3.2.0). All auditory stimuli were presented at an average level of 76 dB (LAeq).

### 2.4 Procedure, task, and experimental design

Figure 1 illustrates the sequence of events in the experiment. A briefly flashing fixation dot signaled the start of each trial for 100 ms (increasing in size from 0.25° to 0.50°). After 1000 ms following the onset of the fixation dot, an audio-visual memory item, consisting of a spatially and temporally synchronized tone and grating, was presented for 250 ms. Tone and grating pairs were randomly chosen on a trial-by-trial basis. Following a 600 ms delay period, an audio-visual impulse stimulus, consisting of a complex tone and a white circle, was presented for 100 ms. The task-irrelevant impulse stimulus was included to revoke potentially activity silent memory traces during the delay (Wolff et al., 2015, 2017, 2020) and assess how their strength relates to subsequently measured neural reinstatement during recall. After a second delay of 700 ms, a probe item was presented for 250 ms. The latter could be either visual (25%), auditory (25%), or audio-visual (i.e., conjunction, 50%). Critically, presenting all probe types equally (i.e., 1/3 of each) would have biased task demands toward segregated feature storage, as participants would have been tested on a unimodal feature in 2/3 of trials and would have retrieved the whole object (conjunction condition) in only 1/3 of trials. Therefore, we deliberately presented single-feature probes in half of the trials, and conjunction probes in the other half. To balance physical stimulation across trials with different probe types, unimodal probes were always presented with a filler; i.e., visual probes were paired with an auditory filler (i.e., pink noise), and auditory probes were paired with a visual filler (i.e., a grey circle). The three types of probes were randomly intermixed throughout the experiment.

**Figure 1.**
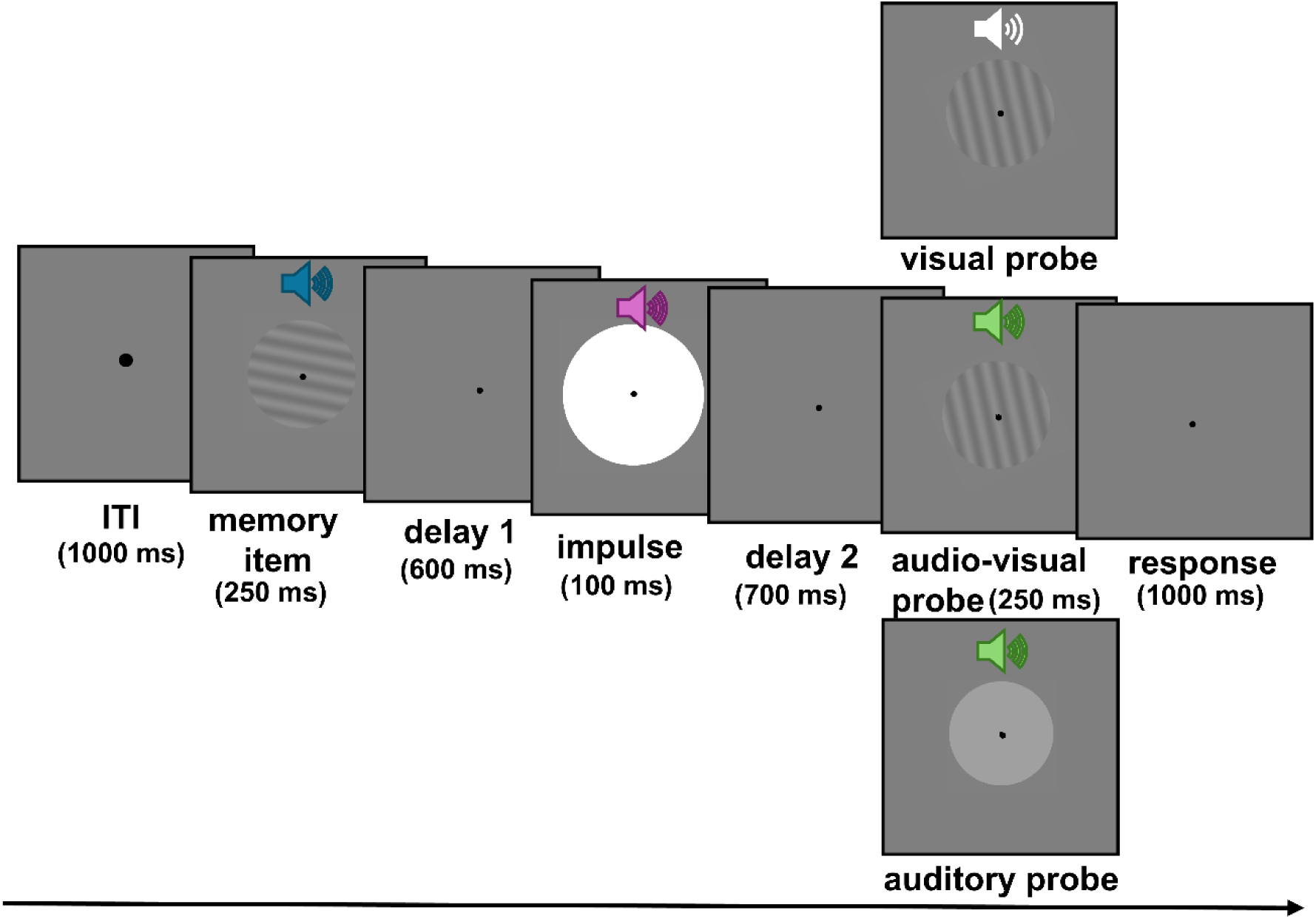
Illustration of the trial sequence. In this delayed match-to-sample task, participants were presented with one audio-visual memory item. After a short delay, a neutral audio-visual impulse stimulus was presented. Following a second delay, participants were presented either with an audio-visual (50%), visual (25%), or auditory (25%) probe item, in a randomized order. Participants indicated via button press whether the probe item matched their memory contents.

Participants indicated whether the probe feature(s) matched the item they were holding in working memory. For conjunction trials, they were instructed that a correct “yes” response required both features to correspond to the previously presented memory item. Participants were not explicitly informed that conjunction trials never included any partial repetitions (e.g., a matching visual feature paired with a novel auditory feature). For unimodal probes, participants were instructed to base their response on the relevant feature and to disregard the fillers.

Accuracy and speed were equally emphasized. Participants responded by using one of two horizontally aligned buttons on a response pad with their index and middle fingers of their dominant hand. The assignment of ‘yes’ and ‘no’ responses to the two buttons was counterbalanced across participants. Responses were permitted at any time from probe onset through the subsequent 1000 ms response period.

In total, the experiment comprised 2304 trials, including 50% trials with conjunction probes and 50% with unimodal probes (i.e., 25% visual, 25% auditory). The trials were divided into 18 mini-blocks of 128 trials each, taking approximately 7 min. At the end of each block, a feedback screen informed participants of their average response time and accuracy for the previous block. All mini-blocks adhered to the same ratio of conjunction and unimodal probe trials. Self-paced breaks were encouraged in between mini-blocks. After half the blocks were completed, participants took an extended 30-minute break, during which they could leave the recording chamber. Before the start of the experiment, participants were once shown the complete set of stimuli to familiarize themselves with the materials and then completed 30 practice trials. Overall, the experimental session took approximately four and a half to five hours.

### 2.5 EEG recording and preprocessing

EEG data were collected using a 64-channel Ag/AgCl electrode cap (BrainCap; Brainvision, Gilching, Germany), with electrodes placed according to the international 10–20 system. Data were recorded at a 1000 Hz sampling rate using a NeurOne Tesla amplifier (Bittium Biosignals Ltd., Kuopio, Finland), with AFz as the ground and FCz as the online reference. Electrode impedances were kept below 20 kΩ.

#### 2.5.1. Preprocessing for time-frequency analyses

For offline processing, continuous EEG data were bandpass filtered between 0.05 and 30 Hz using MATLAB (R2022a) and EEGLAB (Delorme & Makeig, 2004). Noisy channels, with normalized kurtosis exceeding five standard deviations from the mean, were identified and removed using EEGLAB’s *pop_rejchan* function. On average, 4.29 channels were excluded per participant (range = 0–7, *SD* = 1.75) and were subsequently interpolated using spherical spline interpolation. The data were then re-referenced to the average of all channels.

Next, we applied a semi-automatic artifact rejection procedure based on rank-reduced independent component analysis (ICA). First, the continuous EEG data were down-sampled to 200 Hz to facilitate faster ICA decomposition. To increase the proportion of near-dipolar components, the data were additionally high-pass filtered at 1 Hz (Winkler et al., 2015). To further reduce computational demands, only every other trial was retained after segmenting the data into epochs from −1 to 2.9 seconds relative to the onset of the memory item. To optimize the signal-to-noise ratio of the ICA input, an automatic trial rejection procedure (*pop_autorej*) was then applied to remove epochs containing large artifacts or extreme voltage fluctuations (> 1000 µV). This iterative procedure rejected trials exceeding a 5-SD threshold, with a maximum of 5% of trials removed per iteration. ICA was subsequently computed.

The resulting ICA decomposition was subsequently applied to the filtered, re-referenced continuous dataset, which retained its original sampling rate of 1 kHz. The data were then epoched from −1 to 2.9 sec relative to the onset of the memory item and baseline-corrected using a pre-stimulus interval from −200 to 0 ms. Artefactual independent components (ICs) associated with eye movements, cardiac or muscle activity, line noise, channel noise, or other non-neural sources were identified using the EEGLAB plugin *ICLabel* (v1.4; Pion-Tonachini et al., 2019). ICs were rejected if *ICLabel* assigned them a probability of less than 30% of belonging to the *brain* category. On average, 3.38 components (*SD* = 1.16) were rejected per participant. Remaining epochs containing large voltage fluctuations (i.e., exceeding ±150 µV) were excluded using the EEGLAB function *pop_eegthresh*. Trials with premature responses (within 150 ms after probe onset) were discarded, while responses made after 1 second following probe offset were marked as missing. Following this procedure, an average of 534.81 trials (*SD* = 29.30; 92.85%) remained in the auditory condition, 534.11 trials (*SD* = 28.14; 92.73%) in the visual condition, and 1065.78 trials (*SD* = 56.92; 92.52%) in the conjunction condition.

#### 2.5.2. Preprocessing for multivariate pattern analysis (MVPA)

The preprocessing procedure for multivariate pattern analysis followed a similar procedure described above, with minor differences. These differences were introduced to more closely follow the preprocessing scheme for MVPA proposed by Wolff et al. (2020). First, offline continuous EEG data were downsampled to 500 Hz and bandpass filtered between 0.1 and 40 Hz using MATLAB (R2022a) and EEGLAB (Delorme & Makeig, 2004). Noisy channels, with normalized kurtosis exceeding five standard deviations from the mean, were identified and removed using EEGLAB’s *pop_rejchan* function. On average, 4.47 channels were excluded per participant (range = 2–7, *SD* = 1.57), and were subsequently interpolated using spherical spline interpolation. The data were then re-referenced to the average of all channels.

To remove eye-movement artifacts, a semi-automatic procedure based on rank-reduced ICA was used. For this step, data were downsampled to 200 Hz and high-pass filtered at 1 Hz (Winkler et al., 2015) to improve ICA decomposition. To reduce computational load, only every second trial was used after segmenting the data into epochs from −1 to 2.9 seconds relative to memory item onset. Additionally, large-amplitude artifacts were removed using EEGLAB’s *pop_autorej* function, which excluded trials exceeding 1000 μV and iteratively rejected those beyond 5 standard deviations (max 5% per iteration). ICA was computed on this high-pass filtered and cleaned subset of the data. The resulting ICA weights were then applied to the original, filtered, and re-referenced data (with an original sampling rate of 500 Hz). The data were again segmented to create epochs from −1 to 2.9 s relative to memory item onset. Eye-related components were identified using the *ICLabel* plugin (v1.4; Pion-Tonachini et al., 2019) and components with a probability greater than 30% for the “eye” category were removed. On average, 3.28 ICs were excluded per participant (*SD* = 1.14).

Following ICA, remaining trials with voltage fluctuations outside the ±150 μV range were excluded using EEGLAB’s *pop_eegthresh* function. Trials with premature responses (within 150 ms after probe onset) were excluded. Five participants’ datasets were discarded due to extremely noisy data (rejected trials > 25% of all trials). For the remaining subjects (n = 32), an average of 535.65 trials (*SD* = 27.79; 92.58%) remained for the auditory condition, 533.84 trials (*SD* = 28.45; 92.54%) for the visual condition, and 1070.88 trials (*SD* = 53.89; 92.88%) for the conjunction condition.

### 2.6 Multivariate pattern analysis

This analysis aimed to determine whether neural activity during the delay and response periods contained information about the tone frequencies and visual orientations held in working memory. To address this, we adapted the decoding approach developed by Wolff et al. (2020).

To track the temporal dynamics of neural activity patterns, a sliding time window approach was employed that calculated mean-centered voltage patterns within a 100-ms window. Each 100-ms segment was first downsampled by averaging the EEG signal every 10 ms, yielding 10 voltage values per channel. These values were then normalized by subtracting the mean activity of each channel within the window. This normalization step ensured that only temporal fluctuations relative to the window’s baseline were captured. For each trial and each time window position, this procedure yielded a data matrix consisting of 10 time points × 64 channels, which served as the feature space for the eightfold cross-validation decoding procedure, described below. The 100-ms window was then moved forward in 8-ms steps to obtain time-resolved decoding estimates.

For cross-validation, trials were randomly divided into eight folds. Seven folds were used to compute average neural activity patterns for each auditory and visual stimulus (i.e., tone frequencies and orientations in the training set). At this stage, an equal number of trials per stimulus type was ensured by randomly sampling from the available trials. The Mahalanobis distance (De Maesschalck et al., 2000) between the neural activity of each left-out test trial and the averaged activity pattern of each stimulus category was computed, using a covariance matrix estimated from the training data (Ledoit & Wolf, 2004). Since there were eight auditory and eight visual stimulus classes, this procedure yielded eight distance values for each auditory and visual stimulus for each iteration of the cross-validation (i.e., for each left-out trial). This process was repeated 50 times, each with a new random division of trials into folds, and distances were averaged across these repetitions. To facilitate interpretation, the resulting eight distances for each auditory and visual stimulus type were sign-reversed, so that higher values indicate greater similarity between the test and training patterns.

Decoding strength was quantified in two ways: (1) for tone frequencies, a linear regression was conducted between the absolute difference in tone frequency and the corresponding distance value; for tone decoding, the resulting regression slope (i.e., beta value) then served as a measure of decoding accuracy, with positive beta values indicating a systematic increase in neural dissimilarity with increasing tone differences (2) for orientations, cosine-weighted means of distances were computed. For orientation decoding, the cosine-weighted mean reflects decoding accuracy, with higher values indicating increased neural dissimilarity for larger orientation differences.

The analysis was conducted separately for three time periods of interest: from −300 to 700 ms relative to memory item onset, −300 to 700 ms relative to impulse onset, and −300 to 700 ms relative to probe onset. Finally, decoding accuracies were averaged across trials, and the resulting time series were smoothed using a Gaussian kernel with a standard deviation of 16 ms. Importantly, each time point reflected pooled data from the preceding 100 ms (i.e., from −100 to 0 ms relative to that time point). This approach improves the interpretability of decoding onsets, though caution is advised when interpreting decoding offsets (Grootswagers et al., 2017; Wolff et al., 2020).

In particular, we ran decoding analyses to observe whether auditory and visual working memory content were present during item encoding, following the impulse stimulus, and upon probe presentation. Considering our main interest in whether task-irrelevant feature representations were reinstated by unimodal auditory and visual probes, tone frequencies were decoded from visual probes, and orientations were decoded from auditory probes. Additionally, by using the same methodology described here, auditory and visual working memory content were decoded from task-relevant probe conditions. However, for this analysis, only trials during which the probe did not match the memory content were used. This was done to ensure that memory decoding was independent of physically identical probe features (see Supplementary Materials, section 1 for the results of those analyses).

Subsequently, Pearson’s *r* correlations were computed to examine the relationship between decoding strength during the delay and response periods. This analysis tested whether the degree of feature-specific decoding during the delay could predict the extent of neural reinstatement of the task-irrelevant feature upon unimodal probe presentation. Finally, a series of Pearson’s *r* correlations was computed to determine whether impulse-evoked decoding predicted behavioral costs. Specifically, we correlated (1) impulse-evoked orientation decoding with behavioral performance (separately for accuracy and RTs) in auditory probe trials, and (2) impulse-evoked frequency decoding with behavioral performance in visual probe trials.

### 2.7 Time-frequency decomposition

Event-related spectral perturbations (ERSPs; Makeig et al., 2004) were computed using Morlet wavelet convolution via the *pop_newtimef* function in EEGLAB. EEG epochs were convolved with complex Morlet wavelets spanning 4-30 Hz, distributed logarithmically across 52 steps. To optimize the trade-off between temporal and frequency resolution, the number of wavelet cycles increased progressively with frequency, starting at 3 cycles at the lowest frequency and reaching 15 cycles at the highest, with a factor of 0.5.

Following Grandchamp & Delorme (2011), baseline correction was performed using a two-step approach. First, single-trial data was normalized by applying a full-epoch length single-trial correction. Then, a common trial-average baseline was applied, computed over the −400 to −100 ms stimulus interval (relative to memory item onset). This approach has been shown to enhance the signal-to-noise ratio and minimizes contributions from artifactual data (Grandchamp & Delorme, 2011). Only trials with accurate responses were included in this analysis. The resulting ERSP segments extended from −582 to 2482 ms relative to the memory item.

### 2.8 Statistical analyses

Statistical analyses were performed using MATLAB® (R2022a) and JASP (version 0.19.3.0). An alpha level of 0.05 was used to determine statistical significance. Partial eta squared (*η*²p) and Cohen’s d (Lakens, 2013, equation 10) are reported for repeated measures ANOVAs (rmANOVA) and for t-tests, respectively. For rmANOVAs, Greenhouse-Geisser corrected degrees of freedom are reported, in case the sphericity assumption was violated (Mauchly’s test *p* < .05). Bonferroni-Holm correction was applied to correct for multiple comparisons, and adjusted *p*-values are reported as *p_corr_*.

#### 2.8.1 Behavioral analyses

To assess behavioral performance, accuracy (percentage of correct responses) and mean reaction times (RTs) were calculated per condition. Slow responses that occurred after 1 sec following the probe offset were counted as incorrect responses (0.4% of overall trials). To assess performance differences between the three probe conditions, a one-way rmANOVAs was conducted for accuracy and RTs, respectively.

#### 2.8.2 ERSP analyses

To test whether midfrontal theta power was modulated by our experimental manipulation, cluster-based permutation tests were conducted using the FieldTrip toolbox (Version 20.230.422; Oostenveld et al., 2011). Analyses specifically focused on Fz and Cz (Cohen, 2014; Cohen et al., 2008), frequencies in the theta range between 4–7 Hz, and an 800 ms time window during the response period (i.e., - 50 to 750 ms relative to probe onset). For each time-frequency point, a two-sided paired-sample t-test was conducted. Then, time-frequency samples with *p* < 0.025 were selected and clustered based on spectral and temporal adjacency. Each cluster’s statistical value was calculated as the sum of t-values within the cluster, and the maximum cluster statistic was used for determining significance (see Arslan et al., 2025a for more details). To assess statistical significance, a Monte Carlo permutation method was applied. Specifically, trials were randomly assigned a condition label (e.g., visual, auditory, conjunction), and the procedure described above was repeated 1000 times to create a null distribution of cluster statistics. Finally, observed cluster values were considered significant if they exceeded the 99th percentile or fell below the 1^st^ percentile of the null distribution. However, it is important to note that the duration or frequency range of a given cluster should not be interpreted as the exact boundary of the effect. Rather, these tests evaluate whether the overall size of the cluster is unlikely under the null hypothesis (Sassenhagen & Draschkow, 2019). These cluster-based permutation analyses were run to test differences in ERSPs between different probe trials, contrasting visual vs. auditory, visual vs. conjunction, and auditory vs. conjunction trials, and Cohen’s d was used as the effect size measure (Lakens, 2013, equation 6). Effect sizes for significant clusters were estimated using the average over cluster method (Meyer et al., 2021).

#### 2.8.3 Decoding analyses

To test where the time-course of decoding exceeded chance level (i.e., 0), cluster-corrected sign-permutation tests were conducted. First, at each time point, the sign of each participant’s decoding value was randomly flipped with a probability of 50%. This generates a distribution of test statistics under the null hypothesis that decoding values are equal to zero. To control for multiple comparisons across time, the resulting time-resolved test statistics were entered into a cluster-based permutation test, in which temporally adjacent samples exceeding a cluster-forming threshold of *p* < 0.05 were grouped into clusters. Cluster significance was assessed using permutation testing (100,000 permutations). All tests were one-sided (i.e., testing for decoding above chance).

For significant decoding clusters, effect sizes were quantified using an average over cluster approach (Meyer et al., 2021). Specifically, decoding values were first averaged across all time points within a given significant cluster for each participant, yielding a single cluster-level decoding value per participant. Effect sizes were then computed as Cohen’s d (Lakens, 2013, equation 6).

## 3. Results

### 3.1. Behavioral results

Figure 2 shows the behavioral performance for each probe condition. Analyses revealed significant differences in accuracies between probe types, *F*(1.40, 50.37) = 48.12, *p* < 0.001, *η*^2^_p_ = 0.57. Follow-up paired-samples t-tests showed that performance was higher in conjunction trials than in visual (*t*(36) = 10.46, *p_corr_* = 0.002, *d* = 1.72) and auditory probe trials (*t*(36) = 2.67, *p_corr_* = 0.011, *d* = 0.44). Furthermore, accuracy was higher in the auditory compared to the visual probe trials (*t*(36) = 5.71, *p_corr_* = 0.003, *d* = 0.94).

**Figure 2.**
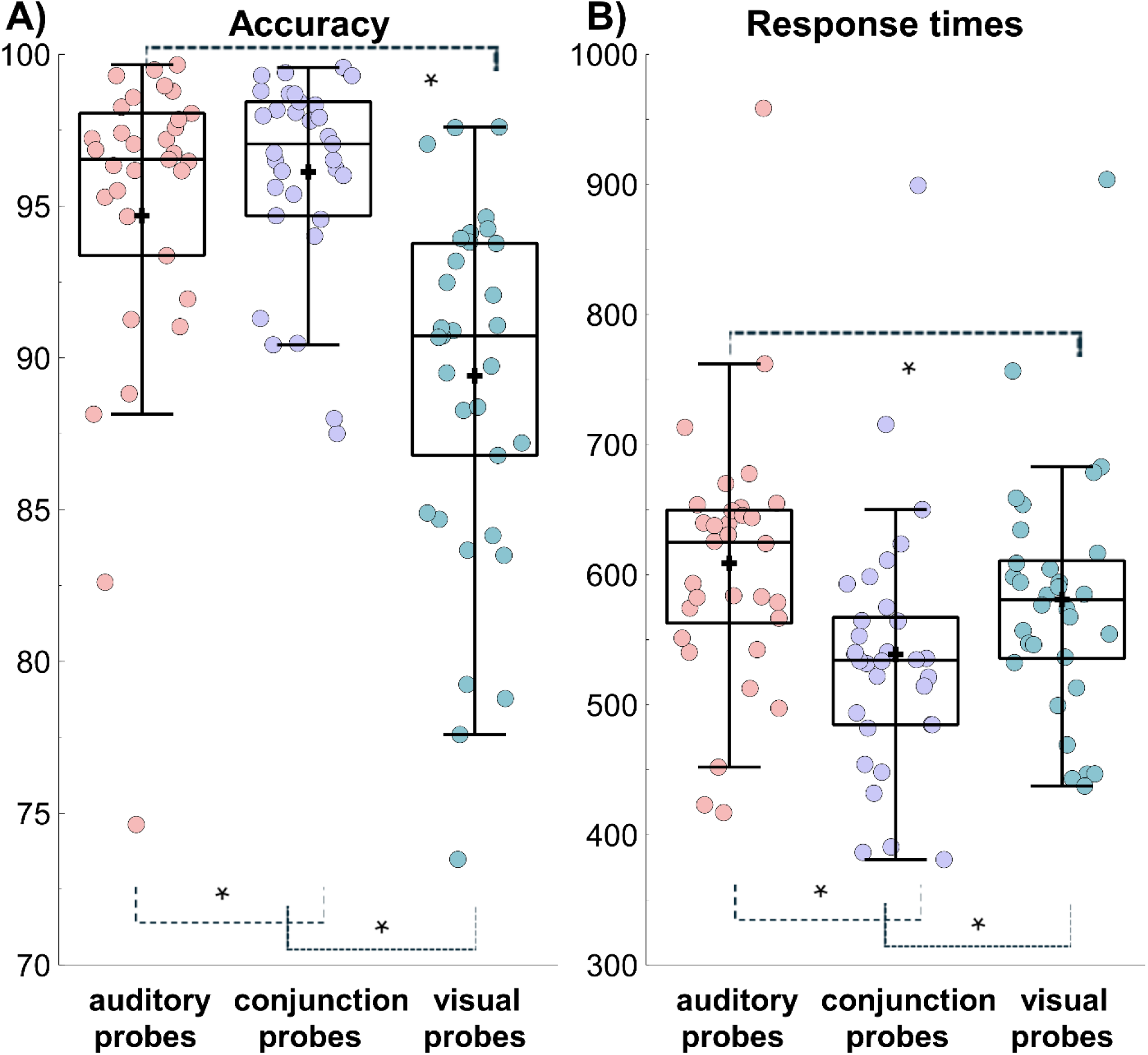
Behavioral performance across probe conditions. (A) The proportion of correct responses; (B) RTs for the conditions. Boxplots show the +/− 1.5 interquartile range and the median. The dots illustrate individual participant averages per condition. A black cross illustrates the condition mean.

Similarly, results revealed significant differences in RTs between probe conditions, *F*(1.44, 51.65) = 89.44, *p* < 0.001, *η*2p = 0.71. Follow-up paired-samples t-tests showed that RTs were faster in the conjunction than in the visual (*t*(36) = 10.31, *p*_corr_ = 0.001, d = 1.7) and auditory probe trials (*t*(36) = 14.44, *p_corr_* = 0.002, *d* = 2.37). Furthermore, RTs were slower in the auditory compared to the visual probe trials (*t*(36) = 4.52, *p_corr_* = 0.003, *d* = 0.74).

Overall, in line with our expectations, behavioral performance improved in conjunction probe trials compared to unimodal probe trials. This is in line with the hypothesized cost of “breaking up” a bound representation.

### 3.2. Decoding tone frequencies and visual orientations

Neural activity patterns contained decodable information about the tone frequency and orientation during memory item presentation and encoding. Specifically, tone frequency and orientation were decodable from 22 – 774 ms (*p* < 0.001, *d* = 2.86, one-sided, corrected) (see Figure 3A) and 70 –758 ms (*p* < 0.001, *d* = 2.35, one-sided, corrected) (see Figure 3D) relative to memory item onset, respectively.

**Figure 3.**
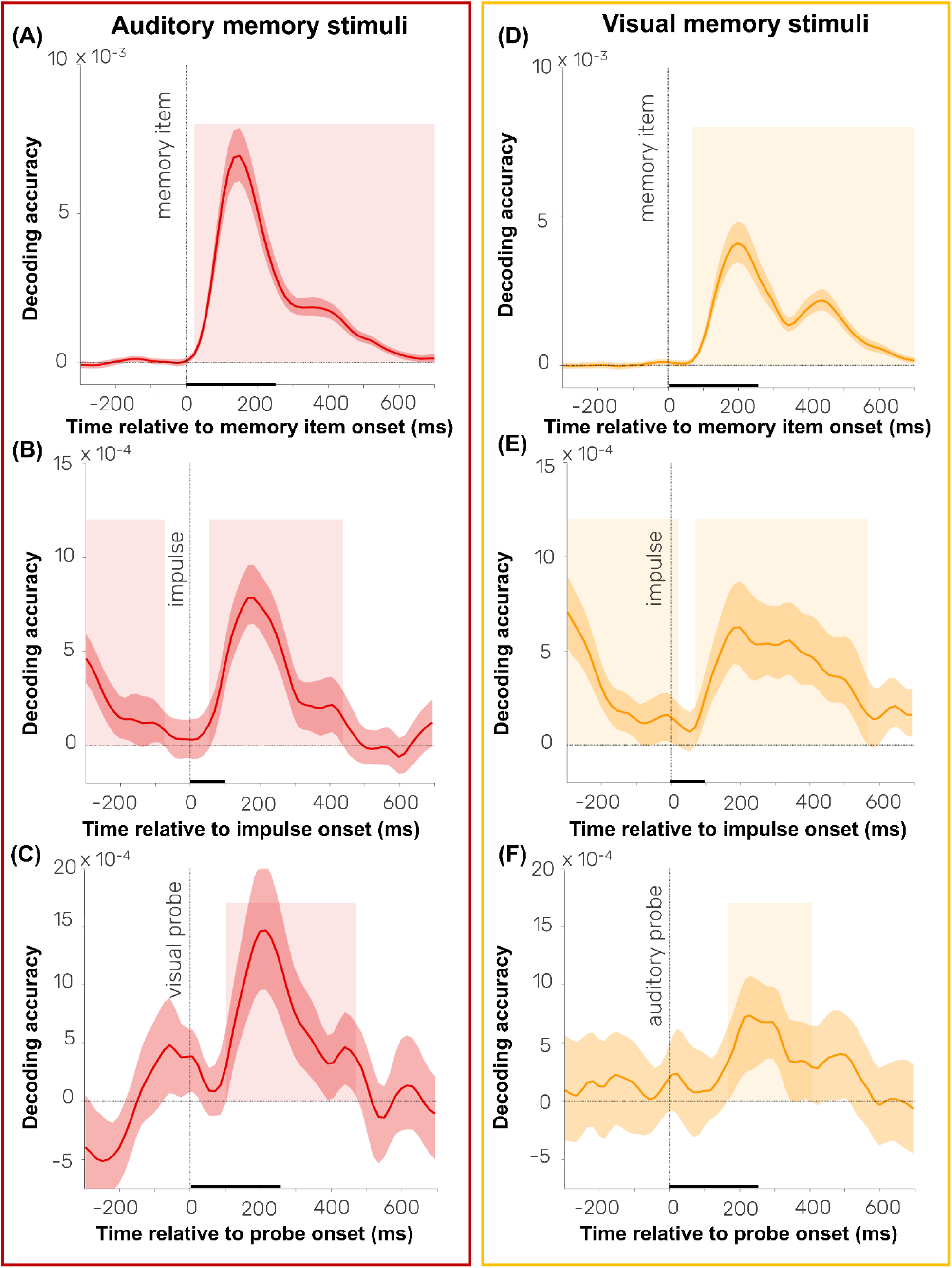
Decoding of auditory and visual memory across different processing stages. Panels (A–C) show decoding time courses for tone frequencies (beta values) upon (A) item encoding, (B) the impulse response, and (C) visual probe presentation. Panels (D–F) display decoding time courses for orientations (cosine-weighted means) upon (D) item encoding, (E) the impulse response, and (F) auditory probe presentation. Shaded regions indicate time points within significant clusters. Black horizontal lines along the time axis indicate the durations of memory items, the impulse stimulus, and the probes. Error shading indicates 95% confidence interval of the mean.

Moreover, feature-specific information in neural activity patterns persisted throughout maintenance and re-emerged upon impulse presentation during the delay period. That is, tone frequency could be decoded both before (−298 to −74 ms, relative to impulse onset, *p* = 0.02, *d* = 0.86, one-sided, corrected) and after the impulse (54 – 438 ms relative to impulse onset, *p* < 0.001, *d* = 1.60, one-sided, corrected) (see Figure 3B). Orientation-decoding revealed a similar pattern, with two significant clusters surrounding the impulse stimulus (−298 to 22 ms relative to impulse onset, *p* < 0.001, *d* = 1.28, one-sided, corrected; and 70 to 566 ms, relative to impulse onset, *p* = 0.008, *d* = 1.11, one-sided, corrected) (see Figure 3E). It should be noted that significant decoding prior to impulse presentation partially overlaps with the significant clusters that were obtained relative to memory item onset.

Importantly, the feature-specific decoding showed significant clusters for tone frequencies upon visual probe presentation (102 – 470 ms relative to probe onset, *p* < 0.001, *d* = 0.92, one-sided, corrected) (see Figure 3C), and for orientations upon auditory probe presentation (166 – 406 ms, relative to probe onset, *p* < 0.001, *d* = 0.73, one-sided, corrected) (see Figure 3F). These results confirm our hypothesis that task-irrelevant features (i.e., tone frequencies in visual probe trials and orientations in auditory probe trials) were reactivated following unimodal probes, consistent with an object-based account. Additional feature-specific decoding of task-relevant features in no-match trials and the comparisons of decoding accuracies across all probe types are discussed in the supplementary materials.

A subsequent analyses aimed at investigating whether the degree of feature-specific decoding during the delay predicts the degree of neural reinstatement of the task-irrelevant feature following unimodal probe presentation. Results showed that the strength of impulse-evoked decoding for orientations was positively correlated with the neural reinstatement of visual features upon conjunction probe presentation (Pearson’s *r* = 0.58, *p* < 0.001). In contrast, the impulse-evoked decoding for orientations was not predictive of orientation-specific decoding during recall upon visual probe presentation (Pearson’s *r* = 0.05, *p* = 0.81) and auditory probes (Pearson’s *r* = 0.34, *p* = 0.06). No significant correlations between impulse-evoked and probe-evoked decoding for tone frequencies were observed, irrespective of the probe condition (all Pearson’s *r* < 0.21, all *p* > 0.25).

Finally, a group of subsequent analyses aimed at investigating whether the extent of feature-specific decoding during the delay predicts the degree of behavioral costs. Results showed that neither the impulse-evoked decoding of orientations predicted the accuracy (Pearson’s *r* = 0.33, *p* = 0.07) or reaction times (Pearson’s *r* = −0.15, *p* = 0.42) observed in auditory probe trials nor the impulse-evoked decoding of frequencies predicted the accuracy (Pearson’s *r* = −0.04, *p* = 0.84) or reaction times (Pearson’s *r* = −0.04, *p* = 0.82) in visual probe trials.

### 3.3. Time-frequency results

Figure 4 shows the time course of midfrontal theta power relative to probe onset and the results of the cluster-based permutation contrasts. As depicted, midfrontal theta power increased more strongly during unimodal than during conjunction probe trials. Specifically, the difference between visual and conjunction probes extends roughly from 50 ms to 750 ms after the probe onset (*p* < 0.001, *d* = 0.22), whereas the difference between auditory and conjunction trials extends roughly from 260 ms to 750 ms after the probe onset (*p* < 0.001, *d* = 0.22). Finally, the difference between visual and auditory probes extends roughly from 40 to 540 ms after the probe onset (*p* < 0.001, *d* = −0.24). Overall, these results confirm that higher cognitive control is required for unimodal than for conjunction probes. Furthermore, the greater cognitive control required by visual than by auditory probes is consistent with lower accuracy with visual probes, suggesting that visual probes were more cognitively demanding, possibly because auditory fillers accompanying visual probes were more salient than visual fillers accompanying auditory probes.

**Figure 4.**
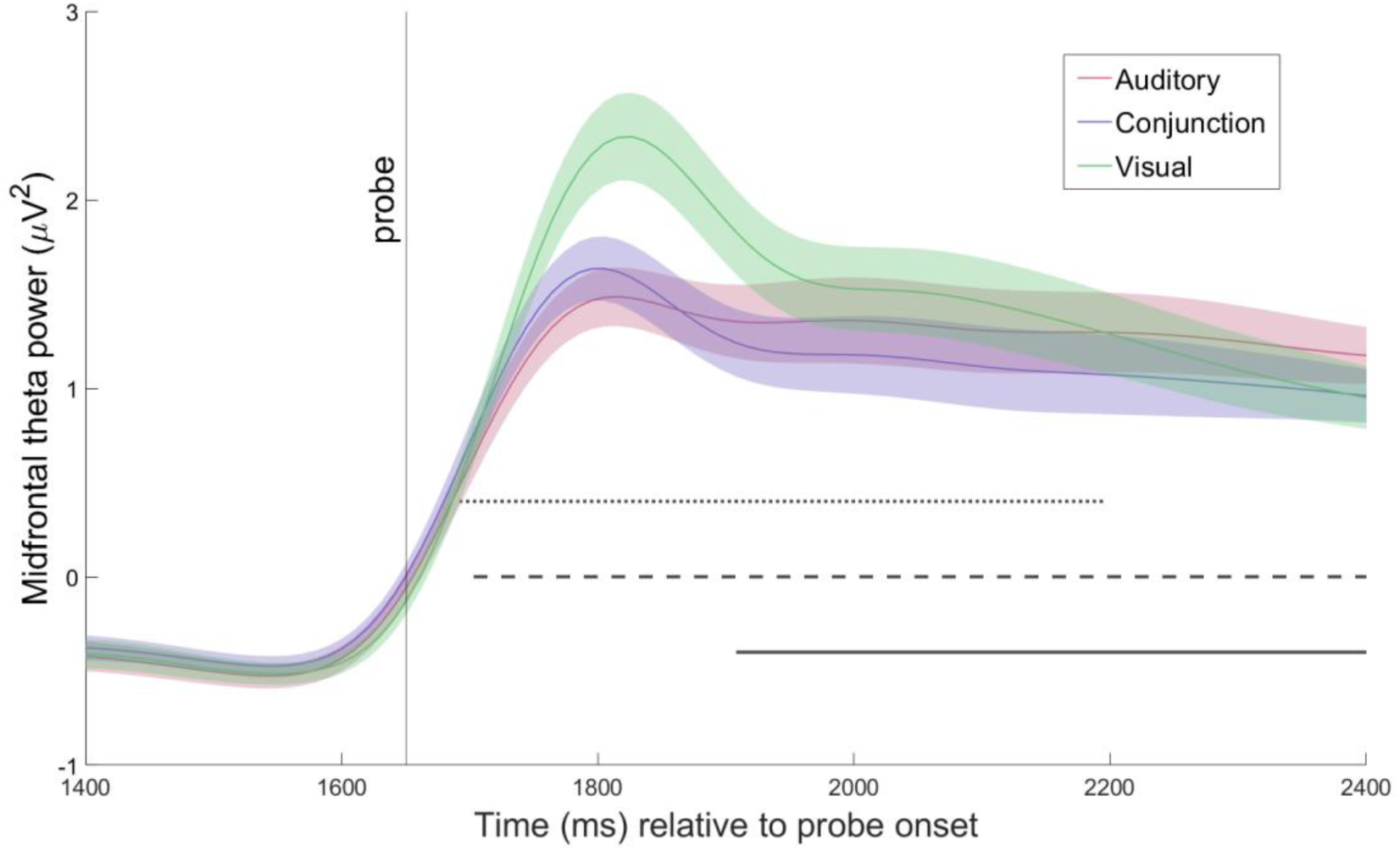
Midfrontal theta power across probe trials and results from cluster-based permutation contrasts. Differences in midfrontal theta oscillatory power were tested with cluster-based permutation tests between visual (green) vs. conjunction (purple), auditory (pink) vs. conjunction, and auditory vs. visual probe contrasts. Results showed significant differences in midfrontal theta power between auditory and conjunction (indicated by a solid line); between visual and conjunction (indicated by a dashed line); and between visual and auditory probes (indicated by a dotted line).

## 4. Discussion

A central question in working memory research concerns whether retroactive attentional selection operates on individual features or integrated object representations. The present study addresses this question from a multisensory perspective by examining whether selectively retrieving a single feature of a cross-modal object in working memory results in the (involuntary) recall of the entire object. Specifically, we ask whether the cross-modal spread of attention across object features is reflected in behavioral costs during single-feature recall and the neural reinstatement of task-irrelevant features. The current study reveals three key findings: (1) recalling a single feature of an audiovisual object incurs a behavioral cost relative to recalling the entire object; (2) this is reflected in neural indices reflecting the reactivation of task-irrelevant features following modality-specific probes, i.e., in line with object-based recall; (3) this reactivation increases cognitive control demands when feature-based prioritization is required. Together, these results offer further insights into the nature of audiovisual object representations in working memory, suggesting that these are stored as bound units.

The behavioral cost, associated with recalling a single feature of an audiovisual object, is in line with prior work showing that task-irrelevant features can impair selective attention to relevant sound features (Maddox & Shinn-Cunningham, 2012). Moreover, it closely mirrors the feature extraction cost previously reported in auditory working memory (Joseph et al., 2015), suggesting that object-based representations can interfere with feature-based prioritization. In the study by Joseph et al. (2015), the behavioral cost was attributed to interference both at encoding and recall. That is, given that the authors cued the task-relevant feature in a block-wise fashion (i.e., participants knew they needed to memorize only the temporal or spectral features of a sound at the time of encoding), interference from the irrelevant feature dimension may add noise to the working memory representation already during encoding. In addition, they showed that the feature extraction cost was greater when the irrelevant feature of the probe item was randomly changed rather than held constant, indicating that the effect could also be partially driven by interference from irrelevant features during retrieval. In contrast, in the present study, the task structure allows us to assume that the cost of extracting one feature dimension arises solely at retrieval, when the probe indicates the to-be-tested feature.

A similar form of incidental retrieval of bound object features is proposed by the event file framework (Hommel, 2004; Kahneman et al., 1992; Treisman & Zhang, 2006), where the repetition of one - but not all - object features is assumed to reactivate the entire event file, including previously co-occuring features and responses. Empirical studies, showing partial repetition costs in an audiovisual context (Zmigrod et al., 2009; Zmigrod & Hommel, 2010) are in line with this theory. Moreover, the current results further resemble findings from episodic memory research, demonstrating sensory reactivation during retrieval (Nyberg et al., 2000; Vaidya et al., 2002; Wheeler et al., 2000). In these studies multisensory binding during encoding gives rise to integrated memory traces that can be spontaneously reinstated when a single component of the original event is probed. Extending these findings to working memory, our results suggest that the concurrent presentation of auditory and visual features facilitated audiovisual binding (Meredith & Stein, 1986; Meredith et al., 1987) and storage of those cross-modal features as bound units. In consequence, they tend to be retrieved in an object-based manner, such that the selection of a single feature dimension automatically reactivates associated, task-irrelevant features.

This reactivation of task-irrelevant object features during feature-selective retrieval imposes greater demands on cognitive control. This was reflected by elevated midfrontal theta activity when participants had to retrieve a single feature. By contrast, midfrontal theta power was reduced following conjunction (relative to unimodal) probes, consistent with reduced regulatory demands when retrieval requirements aligned with object-based representations. The reduced cognitive control demands and better memory performance following conjunction probes align with the previous research showing enhanced memory performance when encoding and retrieval formats match (Blaxton, 1989; Cabeza, 1994; Parks, 2013; Pecher et al., 2025; Roediger & Blaxton, 1987; Zeelenberg, 2005).

The notion of incidental object-based retrieval is further corroborated by our EEG analysis. Using multivariate pattern analysis, we show that auditory *and* visual features were not only represented during the delay period, but also following unimodal probe presentation, when one modality becomes task-irrelevant. Specifically, we observed feature-specific decoding not only in the modality that had to be recalled, but also for its associated, now task-irrelevant modality. That is, memory orientations were represented in the neural activity patterns following auditory probes (Figure 3F), and tones were represented following visual probes (Figure 3C). These findings indicate that probing a single feature incidentally reactivates the entire audiovisual object, including its task-irrelevant components. This pattern of neural reinstatement parallels findings by Printzlau et al. (2022), who showed that incidental color cues presented during maintenance result in the neural reinstatement of associated orientation features. A critical difference to this earlier study, however, lies in the functional relevance of the reactivated object features. In the study by Printzlau et al (2022), as noted by the authors, the reinstated feature remained at least potentially relevant for subsequent recall. That is, although the incidental cues were unpredictive of the later tested item, in 50% of trials, the tested item was congruent with the item that was cued on the incidental task. In contrast, in the present task, neural reinstatement is observed at a time, when the reactivated feature was no longer required and could have, in principle, been removed from working memory. The persistence of decodability of these task-irrelevant features thus suggests that removal of no-longer relevant object features from working memory may not occur instantaneously; rather, those features may be deprioritized in a gradual manner (Arslan et al., 2025b). This contrasts with findings suggesting that rapid removal may be possible under certain conditions, such as for verbal materials cued to be irrelevant shortly after encoding (Oberauer, 2018).

The present study also included an audiovisual impulse (i.e., a “ping”) in the delay period, to perturb potentially activity-silent representations. Previous studies had shown that trial-wise variability in the decoding of the impulse response predicted accuracy in subsequent working memory guided responses (Wolff et al., 2017). More specifically, earlier reports showed that higher impulse-evoked decoding for the uncued item were associated with a drop in performance for reporting the cued item. Here, we additionally tested the hypothesis that decoding strength during the delay (i.e., following the presentation of the ping) would predict the size of the behavioral extraction cost and the degree of neural reinstatement following unimodal probe presentation. However, the only significant association found was between the strength of impulse-evoked decoding for orientations and the neural reinstatement of visual features following conjunction probe presentation. In contrast, the strength of impulse-evoked decoding did not predict the behavioral cost or degree of neural reinstatement in unimodal probe trials. This suggests that impulse-evoked decoding indexes the availability of feature-specific representations during maintenance, but does not capture the strength of binding between features that drives subsequent behavioral costs.

At first glance, the observed behavioral extraction cost appears to contrast with evidence showing performance benefits of retro-cues, which allow attention to be directed toward task-relevant features (Hajonides, et al., 2020; Heuer et al., 2016; Heuer & Schubö, 2016; Park et al., 2017). One possible reason for this discrepancy is that retro-cues delivered during maintenance are subsequently followed by an extended delay-period. This allows time for the dynamic modulation of memory representations prior to test, including the strengthening of relevant features (Li et al., 2023) and the attenuation of irrelevant ones (Klatt et al., 2020; Rösner et al., 2020). In the present study, by contrast, probes simultaneously directed attention to the task-relevant feature dimension(s) (akin to a retro-cue) and acted as the test item, against which working memory contents were evaluated. By design, it is therefore difficult to differentiate between prioritization and retrieval in the present study. We would predict that the behavioral costs may be attenuated if a valid retro-cue had indicated the to-be-tested modality prior to presenting the test item. In the present study, by contrast, probes simultaneously directed attention to the task-relevant feature dimension(s) (akin to a retro-cue) and acted as the test item, against which working memory contents were evaluated. By design, it is therefore difficult to differentiate between prioritization and retrieval in the present study. We would predict that the behavioral costs may be attenuated if a valid retro-cue had indicated the to-be-tested modality prior to presenting the test item.

Thus, the present results do not argue against effective feature-based selection. Rather, they align with the notion of ‘hierarchical feature bundles’ as the unit of representations (Brady et al., 2011). According to this view, working memory representations contain both an integrated item-level at the top and individual low-level feature representations at the bottom. The two levels are thought to be at least partially independent, thus, accounting for findings such as the independent forgetting of individual features (Fougnie & Alvarez, 2011). Previous work has shown that attention can operate at both levels (Hajonides et al., 2020) and that the degree of dependence between object features can also be influenced by task demands (Sone et al., 2021). While conjunction and single-feature probes were equally likely in the present study, it remains possible that bottom-up factors, such as the temporal and spatial alignment of audio-visual features, promoted a stronger reliance on object-based storage. Yet, participants were still able to perform the task well-above chance-level, even for feature-selective probes. Thus, the data are not compatible with a strong object-based account, where the sole unit of attentional selection is integrated objects (Lin et al., 2020; Peters et al., 2015, 2021).

Overall, this interpretation converges with our previous work, providing insights into the representational structure of audiovisual working memory (Arslan, et al., 2025a; 2025b). Specifically, we showed that task-irrelevant features are automatically bound into a unified object representation, even when valid pre-cues direct attention to a specific modality (Arslan, et al., 2025a; 2025b). Yet, during maintenance, feature-based attention was evident in modality-specific alpha power modulations (Arslan et al., 2025a). Consistent with this, using representational similarity analysis, a subsequent study showed a gradual shift of activity patterns towards the attended unisensory modality, without fully diverging from a conjunction code (Arslan et al., 2025b). This corroborates the notion that attention selectively prioritizes specific features within bound cross-modal objects. Taken together, this body of work clarifies how hierarchical object-representations and feature-based attentional control jointly shape audiovisual working memory.

Lastly, these findings raise the question of how such incidental reactivation of object-features from partial input may be implemented at the neural level. A candidate mechanism is postulated by models of attention in working memory, which propose that conjunctive neurons encode combinations of features and bind them into stable object representations (Lansner et al., 2013; Manohar et al., 2019). These conjunctive units are thought to remain intact even during activity-silent periods, such that presenting a single feature of a previously encoded object can reactivate the entire conjunctive representation via auto-associative pattern completion. Through this mechanism, associated but task-irrelevant features may be retrieved into an attended state despite not being directly tested.

In conclusion, both behavioral and EEG results suggest that attentional selection of a single feature during retrieval spillover to irrelevant features of the same audiovisual object. Importantly, rather than reflecting a failure of feature-based prioritization, our findings suggest that such feature-based prioritization is cognitively costly because task-irrelevant features are automatically reinstated as part of bound cross-modal object representations. Moreover, our current findings suggest that selectively attending to a single feature within a bound audiovisual object may come at a cost, as it requires breaking up the bound unit or switching between levels of a hierarchically organized representational code. Taken together, the present findings provide a strong test of the object-based account under conditions where the features belong to traditionally distinct sensory modalities (i.e., vision and audition). This challenges the prevailing approach to studying working memory in which the storage of unisensory auditory and visual features is thought to rely on separate storage systems. We provide direct evidence that even semantically unrelated audiovisual features form bound cross-modal objects that can serve as functional units of storage and selection.

## Data and Code Availability

Behavioral and processed EEG data, as well as the analysis scripts and the code supporting the results, are available at the following link: https://osf.io/c4khb. Raw EEG data will be shared upon manuscript acceptance.

## CRediT statement

Conceptualization: CA (lead), LIK (support), DS (support), and SG (support); Methodology: CA (lead), LIK (lead), DS (support); Formal analysis: CA (lead), LIK (support); Writing – Original Draft: CA (lead), LIK (support); Writing – Review & Editing: CA (lead), LIK (lead), DS (support), and SG (support); Visualization: CA; Investigation: CA; data curation: CA; Supervision: LIK; Project administration: LIK (lead), CA (support); Resources: EW.

## Acknowledgements

The authors would like to thank Tobias Blanke for programming the experiment and Mareike Vienken, Clara Weitzenböck, Rama Alnabulsy, Glenn Weilandt and Jane Westedt for their support in collecting the data.

