## Supplementary material for "Neural reinstatement of features in audiovisual working memory indicates object-based retrieval"

### Author Note

### Correspondence

Ceren Arslan, Leibniz Research Centre for Working Environment and Human Factors, Ardeystraße 67, 44139 Dortmund, Germany.

### Supplementary materials

#### 1. Decoding of task-relevant memory features following probe presentation

Supplementary Figure 1 shows the time course of decoding accuracies for task-relevant memory features following probe presentation. In unimodal probe trials, the probe modality indicated which feature dimension had to be recalled, rendering the other feature task-irrelevant at that time. Accordingly, we decoded tone frequency following auditory probes and orientation following visual probes. This supplements the analyses in the main manuscript, demonstrating the decodability of the task-irrelevant features.

To avoid artificial inflation of decoding accuracy due to the presentation of probe stimuli, which, in match trials, contained the same feature we were trying to decode, these analyses included only trials with probes that did not match the memory content (see the 2.6. Multivariate pattern analysis section).

Feature-specific decoding showed significant clusters for frequencies upon auditory probes (182– 342 ms, relative to probe onset, $p$ = 0.02, $d$ = 0.46, one-sided, corrected). Orientation decoding following visual probes revealed two analogous clusters, albeit they fell short of the significance threshold (246 – 342 ms relative to probe onset, $p$ = 0.07, $d$ = 0.57, one-sided, corrected; and 518 – 614 ms, relative to probe onset, $p$ = 0.08, $d$ = 0.43, one-sided, corrected). This absence of statistically significant decoding of task-relevant orientations likely reflects limited statistical power due to the lower number of available trials.

As a robustness check, we thus provide an additional decoding analysis below, including both match- and no-match trials (see the following section).

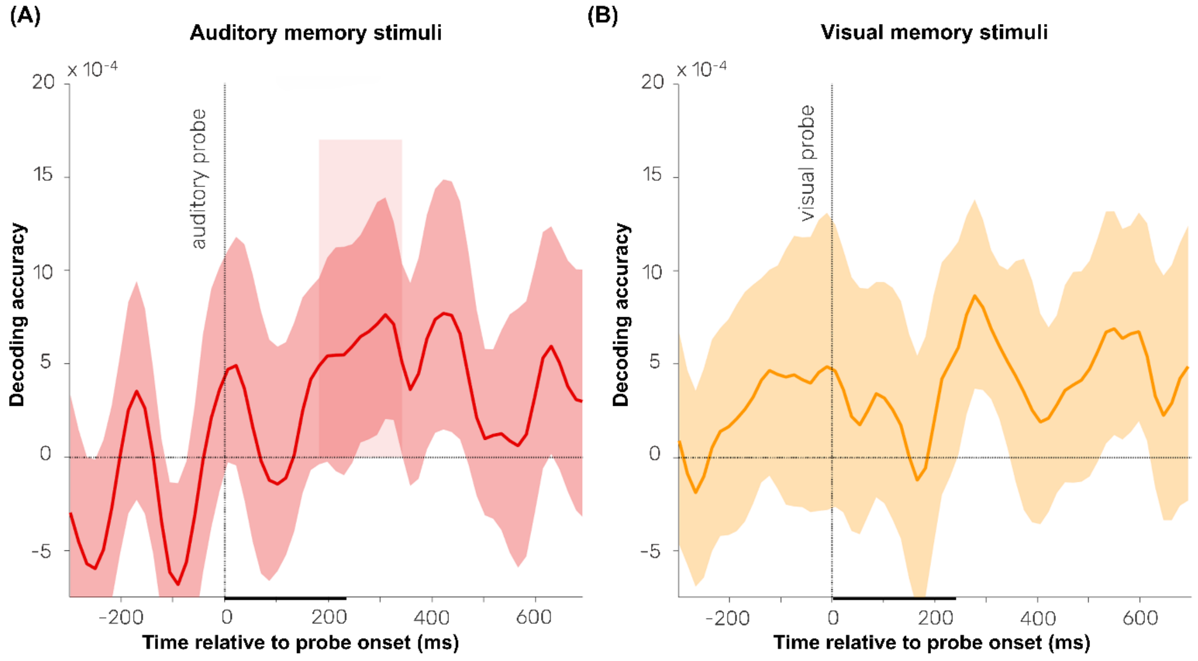


**Supplementary Figure 1. Decoding of auditory and visual working memory content from across task-relevant probes.** Time courses of decoding accuracies for (A) tone frequencies (i.e., beta values) were shown upon auditory probes, and for (B) orientations (i.e., cosine-weighted means) were shown upon visual probes. Shaded regions indicate time points within significant clusters. Black horizontal lines along the time axis indicate the durations of probes. Error shading indicates 95% confidence interval of the mean. These analyses included only 'no-match' trials.

Supplementary Figure 2 shows the time course of decoding accuracies of tone frequencies and orientations upon auditory, visual, and audiovisual (conjunction) probes, when all trials were included. Significant frequency decoding emerged upon auditory (6 – 545 ms, relative to probe onset $p < 0.001$, $d =$ 0.96, one-sided, corrected), visual (102 – 470 ms relative to probe onset, $p < 0.001$, $d = 0.92$, one-sided, corrected) and conjunction probes (54 – 502 ms relative to probe onset, $p < 0.001$, $d = 1.45$, one-sided, corrected). Similarly, significant orientation decoding emerged upon visual (118 – 342 ms relative to probe onset $p < 0.001$, $d = 1.11$, one-sided, corrected; and between 438 – 694 ms relative to probe onset, $p = 0.004$, $d = 0.88$, one-sided, corrected), auditory (166 – 406 ms, relative to probe onset, $p < 0.001$, $d =$ 0.73, one-sided, corrected) and conjunction probes (102 – 662 ms relative to probe onset, $p < 0.001$, $d = 1.70$, one-sided, corrected).

Together, these findings demonstrate that task-relevant features can be robustly decoded during the retrieval phase, when statistical power and signal-to-noise ratio are sufficiently high. However, as noted above, some caution is warranted, as the inclusion of match-trials, in which the probe features correspond to the memorized features, may have inflated decoding performance for task-relevant features via stimulus-driven effects.

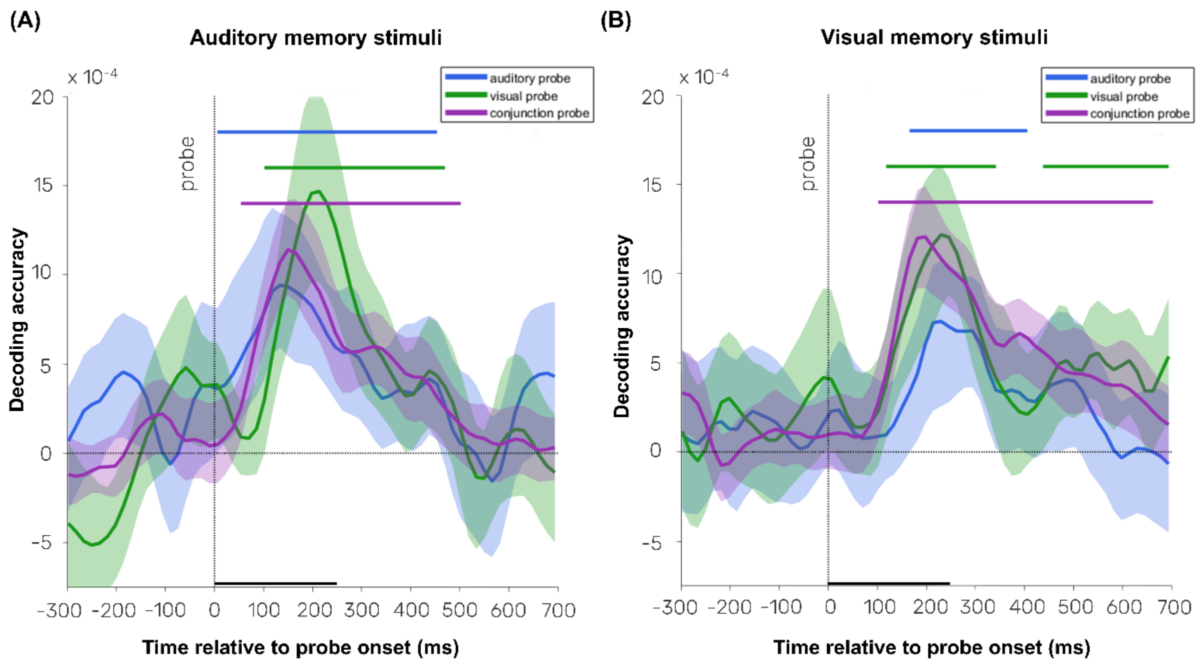


**Supplementary Figure 2. Decoding of auditory and visual working memory content from across different probe types.** Time courses of decoding accuracies are shown for tone frequencies (beta values) and orientations (cosine-weighted means) during and following auditory, visual, and conjunction probes. Colored horizontal lines indicate time points in significant clusters: blue for auditory probes, green for visual probes, and purple for conjunction probes, with each line representing significance for both (A) tone frequency and (B) orientation decoding, respectively. Black horizontal lines along the time axis indicate the durations of probes. Error shading indicates 95% confidence interval of the mean. These analyses included both 'match' and 'no-match' trials.
